# Undersampling techniques for large datasets

**DOI:** 10.1101/2025.09.20.677527

**Authors:** Lexin Chen, Ramón Alain Miranda-Quintana

## Abstract

DNA-Encoded Libraries allow for an efficient approach to synthesize and screen billions of small molecules against a target of interest. With more real-world binding data, this can improve training of machine learning models. However, one key challenge in DELs is the severe imbalances between the classes, in other words, there are much more inactive than active compounds against any given target. This can heavily skew the training process. In this study, we explore different undersampling strategies for the majority class. These different techniques are benchmarked against random selection and prototyped on two different DEL datasets with three different machine learning models. Overall, the `max_sim` strategy shows the best scores, and the general pipeline is implemented in the DELight package.

## Introduction

DNA-Encoded Libraries (DEL) have transformed early drug discovery and how compounds are being screened. Traditionally, high-throughput experiments (HTE) screen tens to hundreds of thousands of compounds each time.^1^ With the chemical space expanding every year, this provided a lot of training data for machine learning models, but screening compounds experimentally would be more costly using HTE.^2,3^ Therefore, DELs have emerged as a key player to be able to screen up to billions of compounds simultaneously, drastically accelerating the early drug discovery timeline.^4,5^ With this key advantage, it also brings along some other things to consider, including the potential for severe class imbalance, with results being inevitably noisy and skewed towards the inactive class.

Usually, the inactive label is more representative in the dataset than the active labels, and this can be problematic for a machine learning model because it will favor false negatives since it sees negatives in the training set more often.^6^ In addition, this can be a problem for the training because most standard loss functions, such as cross-entropy, do not account for class imbalance and will favor the majority class.^7^ Typical ways to mitigate this issue include as undersampling inactives, oversampling actives, SMOTE, or a custom loss function.

Undersampling might be the most intuitive out of these strategies because it simply means downsizing the inactive to match cardinality of the active population.^8–10^ Then the immediate question is which inactives do we choose? The most obvious solution is random undersampling, which randomly selects the same number of inactive as active; however, this can create problems, such as reproducibility.^10^ The second method is cluster-based under-sampling. The idea is to set *k*, the number of clusters, to equal the size of the minority class. Then only the centroids of each cluster will be kept for further analysis. ^11^ This approach has one key disadvantage, in that the centroid is a synthetic point. So, to circumvent this issue, the closest point to the centroid for each cluster will be kept for further analysis, which will solve the synthetic data issue, but pairwise comparisons can be costly. Additionally, the quality of the clusters can diminish the training model if it is not done with care, not to mention the extensive cost of traditional clustering strategies. The third method is NearMiss, which selects the majority class that neighbors the minority class.^12^ The advantage of this approach is that the model to learn the frontier between the majority and minority classes, so it can distinguish between the “tricky” inactives and actives. An issue with this is that it potentially misses important inactives far from active ones that define the differences between the majority and minority classes. The fourth method is the Tomek link,^13^ which aims to draw a clear boundary between the majority and minority classes by removing noise or ambiguous samples near the minority class. The advantage is that this will clarify the distinction between majority and minority classes, but it can also be ineffective if a large portion of the majority and minority classes overlap, because it will not learn the decision boundary in tricky samples.

Another way to balance majority and minority classes is through oversampling, which would entail increasing the size of the minority class to match the size of the majority. One of these ways is to duplicate the minority class samples.^14–18^ Although this is highly efficient and simple, it introduces plenty of artificial points that can make the training model prone to overfitting. Another way is through SMOTE (Synthetic Minority Oversampling Technique).^19,20^ It works by finding the *k*-nearest minority class neighbors and randomly picking one neighbor and creating a new synthetic point along the line between them in the feature space. For example, if you have a minority point at (2,3) and (4,5), SMOTE will make a new point in (3,4). Although this point is not a duplicate and can reduce overfitting, it is a synthetic point, so it is chemically meaningless, and it can introduce noise if the minority class is sparse.

To tackle the class imbalance problem, one approach is the weighted loss function, which would impose an inversely proportional weight to the frequency of that class. ^7,21^ It would work best if the user cares about recall and precision for one class over the other. For example, in the case of fraud detection, there is a severe imbalance between fraud and non-fraud transactions, and the cost of false negatives is much higher than false positives. This can be implemented by using a weighted loss function that assigns a significantly higher penalty to misclassifying the positive (fraud) class, thereby forcing the model to prioritize the reduction of false negatives.^21,22^ It can also raise challenges of how to tune weights properly for classes, and class imbalance problems can still exist even with the punishment system.

For this paper, we are investigating new ways to perform undersampling, as it is the most conventional way of handling class imbalances, and how it affects ML models and different datasets. Despite the advantages of the previously discussed strategies, undersampling does not introduce artificially created points, which we consider to be a key drawback of other methods. The code is available as a GitHub repository at https://github.com/mqcomplab/DELight.

## Materials and Methods

### Data

We used the Aircheck WDR91 and WDR12 datasets https://aircheck.ai/datasets, in partnership with HitGen.^23,24^ WDR91 contains 330K entries, and WDR12 contains 140K entries. Each datapoint has a binary classification label for observed enrichment (0 = not enriched, 1 = enriched). Fingerprints (FP) were pre-generated using the RDKit^25^ graph fingerprint with 2048 bits. These RDKit fingerprints and enrichment labels were used for clustering, undersampling, and model training. The workflow (illustrated in Fig. 1 for the DEL is to first split the training/testing data into 80/20. We paid particular attention to making sure the percentage of actives in the testing data is within 2% of the whole dataset. The data has been split randomly three times. Each split will independently go through undersampling and model training.

**Figure 1:**
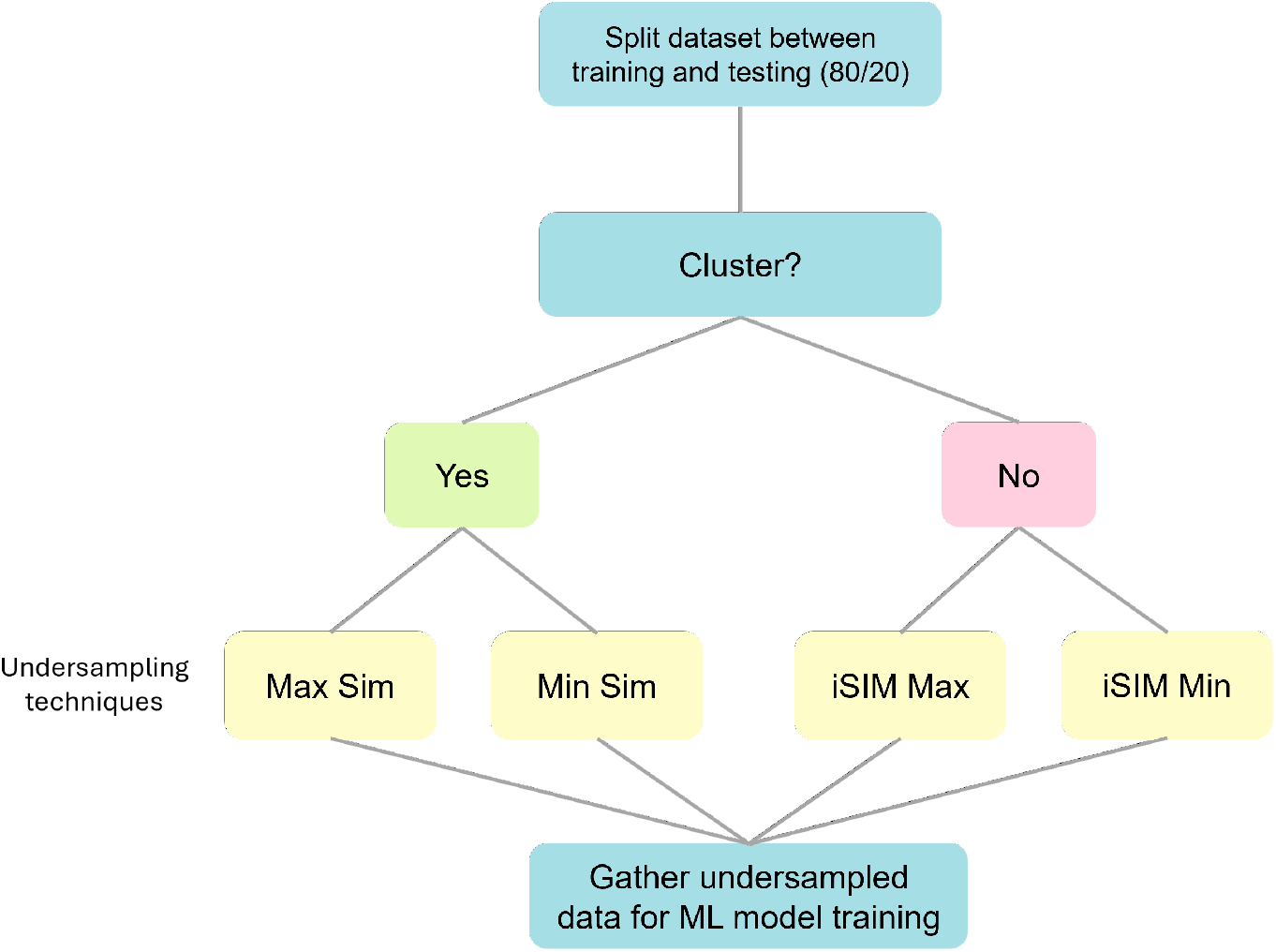
Flow chart for the DEL undersampling workflow.

### Undersampling

Of the four new undersampling techniques explored, two require clustering, and two do not. For the ones that do, it is clustered using BitBIRCH^26^ with BFs refinement,^27^ as a way to ensure that this pipeline is applicable to very large datasets. BFs refinement is a second clustering step after BitBirch, and refinement is needed because BitBirch, forming clusters using a radial threshold, tends to result in one large main cluster with many smaller clusters. Therefore, BFs’ refinement aims to improve the quality of the final assignment. How it works is first, all the members of the big clusters will be broken into individual clusters, and these individual clusters will be reclustered with the other *n −* 1 clusters formed from the first clustering step. BitBIRCH Clustering begins with a threshold of 0.9, and BFs refinement uses the same threshold. If there is a singleton that belongs in the actives, then all the clusters with ten or fewer members will go through another round of clustering at a lower threshold, 0.1 less than the previous. This will continue until there are no more actives as singletons. This clustering step is optional, with our two methods, Max Sim and Min Sim, demanding it a clustering step, but iSIM Max and iSIM Min do not. Max Sim (pseudocode in Alg. 1) aims to sample clusters with both active and inactive molecules so that the model can have a better idea of the decision boundary between the two classes. However, Min Sim (pseudocode in Alg. 2) aims to sample clusters that only have inactives, so there is a clear separation between the two classes for the training model. For iSIM Max and iSIM Min, we look at the similarity using instant similarity (iSIM)^28^ between all the actives and inactives, as a way to speedup the process of navigating through the library. No clustering is needed for this technique. iSIM Max will take the inactives that are most similar to the actives, and iSIM Min will take the inactives that are least similar to the actives.

#### Algorithm 1 max_sim

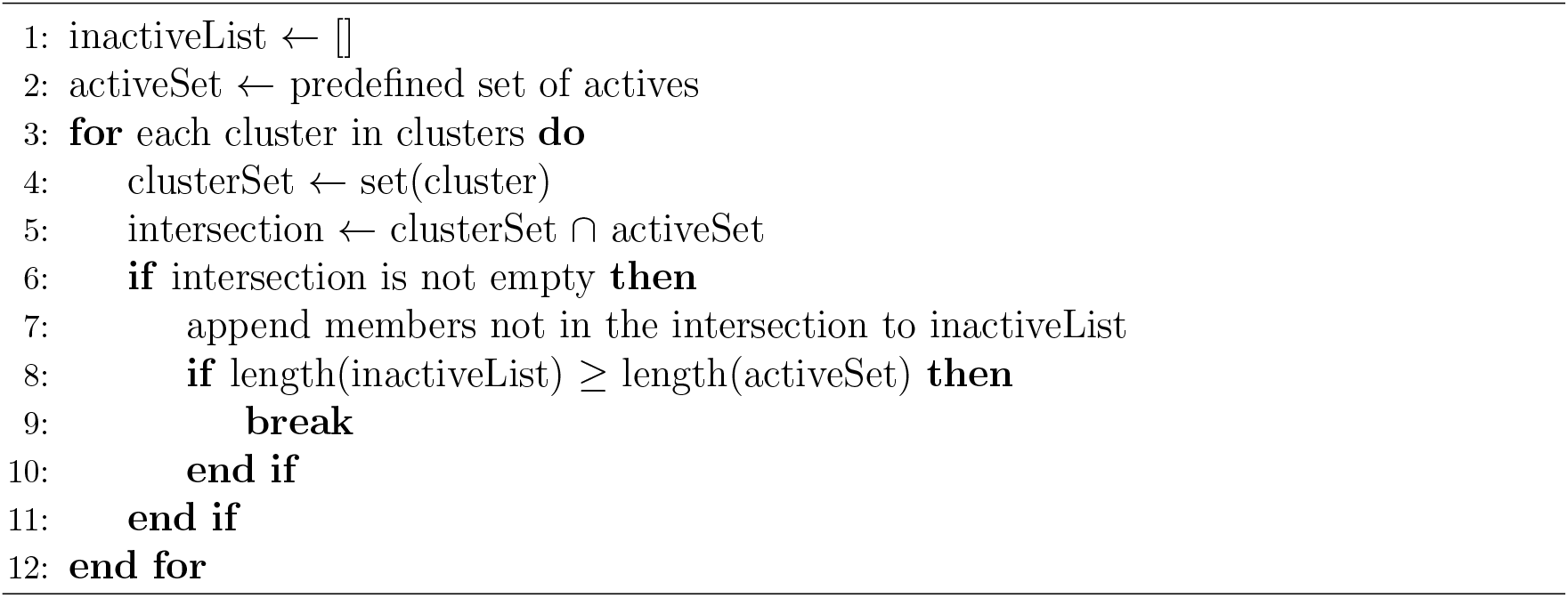

#### Algorithm 2 min_sim

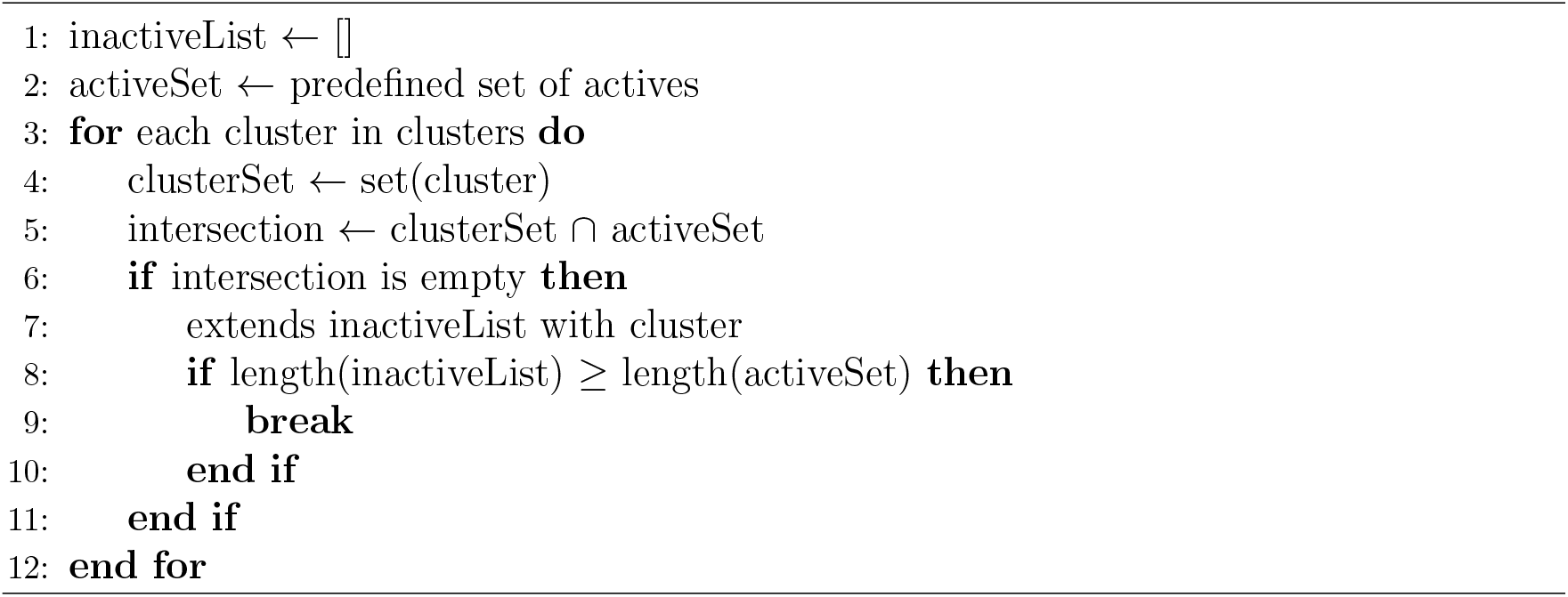

### Machine Learning Models

The logistic regression and random forest models were implemented using scikit-learn with default settings. No feature scaling was applied to these models. For the Multilayer Perceptron, input features were normalized using StandardScaler. The scaler was fit on the training set and then applied to both the training and test sets. The model was built using Keras with two hidden layers (128 and 64 units) using ReLU activation, and a single output neuron with sigmoid activation. It was compiled with the Adam optimizer and binary cross-entropy loss, and trained for 20 epochs with a batch size of 32. 10% of the training data was used for validation. All models were trained and tested on the same dataset for consistent comparison.

## Results

Accuracy calculates how close the predicted labels are to the true labels. A higher score means a closer match to true labels. However, it is important to be wary of this number as higher scores do not always mean a better model. An overfitted model can also have a high score because the model is memorizing the data. In addition, sometimes the cost of error between the classes is not equal. For example, in the case of fraud detection, a false negative is much more malignant than a false positive. Therefore, we have other classification metrics to assess how well the model distinguishes the classes.

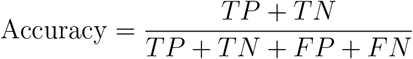

Precision measures the accuracy of positive instances, indicating how many predicted positives are actually correctly labeled. This is critical for identifying false positives, and higher scores would result in fewer false positives.

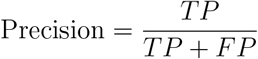

Recall, also known as sensitivity or true positive rate, measures how well the model correctly identifies all the positive instances in the dataset. This is critical for identifying false negatives, and a higher score would mean the model does a good job at recognizing the positive instances and minimizing the number of false negatives.

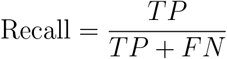

**Figure 2:**
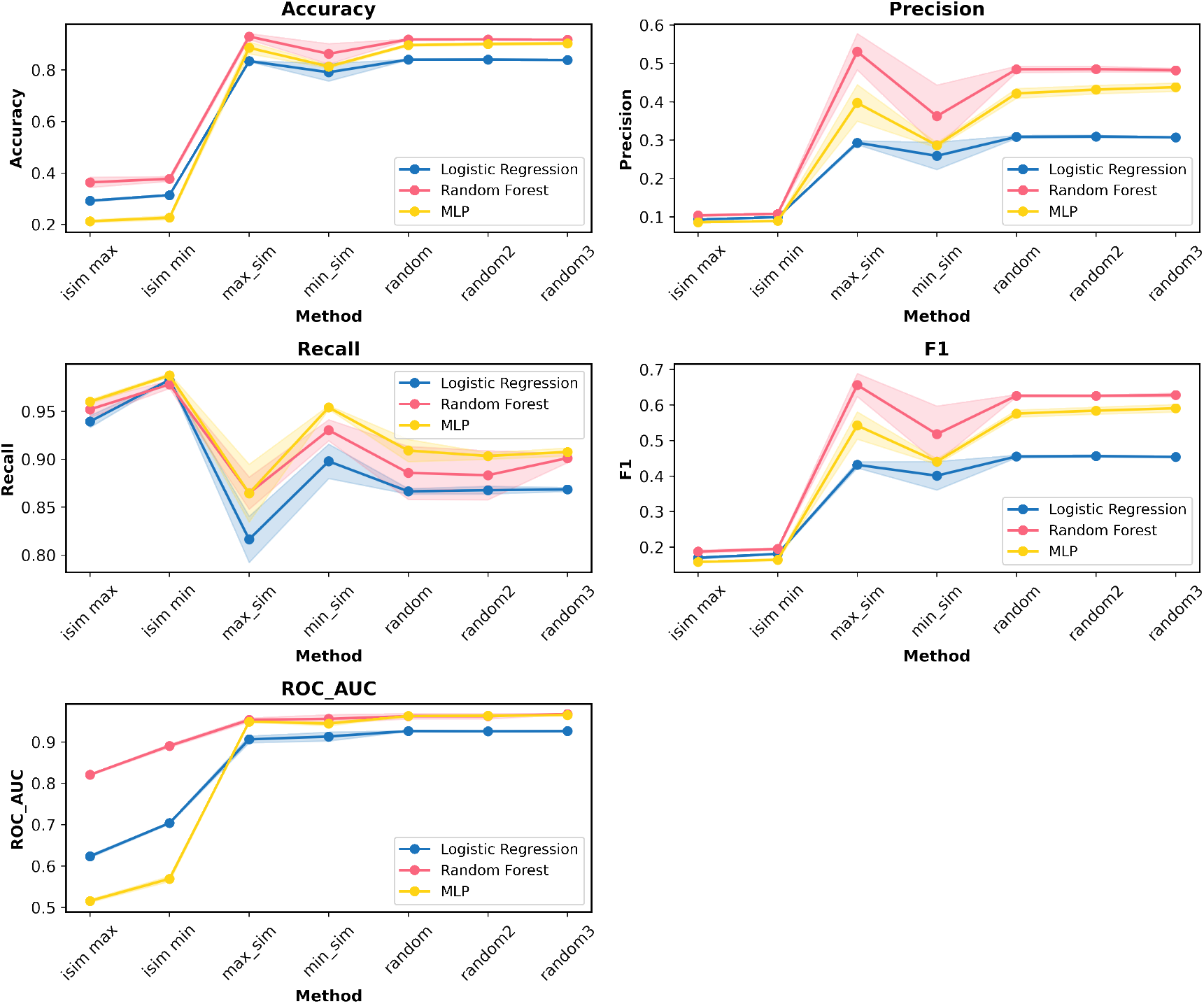
The line graph plots the average score for each metric across the three data splits; the shaded region captures the standard deviation of that metric across the three data splits. This is done for the WDR91 systems.

F1 score uses the harmonic mean of precision and recall. These two metrics have different goals and often require opposite strategies to optimize. To improve precision, you need stricter rules for what counts as a positive prediction—this helps reduce false positives. On the other hand, improving recall means loosening those rules to catch more actual positives, which helps reduce false negatives. Because of this, F1 penalizes the model if either precision or recall is too low. It helps balance the trade-off between the two. This is especially useful in class-imbalanced datasets, where accuracy alone can be misleading. The F1 score gives better insight into whether the model is favoring one class too much or struggling to detect the minority class.

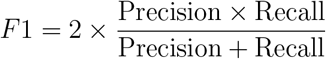

**Figure 3:**
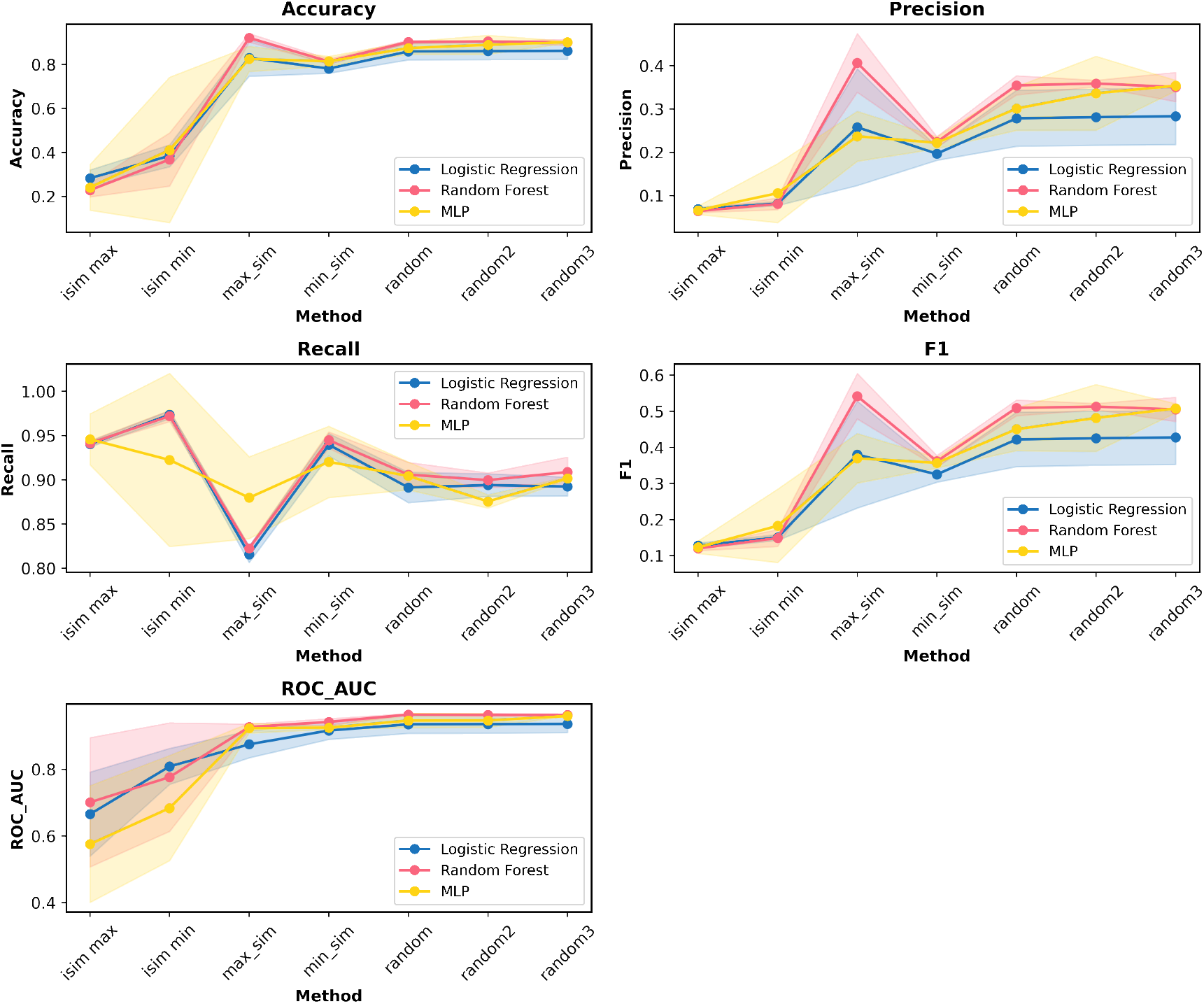
The line graph plots the average score for each metric across the three data splits; the shaded region captures the standard deviation of that metric across the three data splits. This is done for the WDR12 systems.

ROC AUC^29^ measures how well a binary classifier distinguishes between the two classes.

The ROC curve plots the true positive rate (recall) versus the false positive rate at different classification thresholds. The area under the curve (AUC) ranges from 0 to 1, where 0.5 indicates random guessing and 1 indicates perfect prediction. This is not particularly useful for a class-imbalanced dataset, as ROC AUC will be heavily influenced toward the majority class.

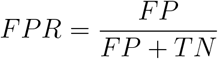

For WDR91, the tradeoff between precision and recall is evident, and they are almost an inverse pattern to one another. In this case, Max Sim performs the best out of all the proposed undersampling techniques. iSIM Max and iSIM Min perform the worst since no clustering is done. This shows that clustering does play a critical role in identifying some of the decision boundaries and exposes the model to a diverse variety of inactives. Seeing from the high recall score, the criteria for defining a compound as positive is very loose, also indicating that it might be labeling most of the molecules as positive because the sample of inactive is too similar to each other, while there is more variety in the active, so the system is more likely to assign the testing compound as active. As shown in the low F1 score, a high recall score does not mean it is a good model if the precision score is low. In this case, F1 might be a better indicator than ROC AUC if the model is leaning more toward one label over the other. In terms of the two techniques that require a clustering technique (Max Sim and Min Sim), they both perform relatively well, and the results are close to the random benchmark scores. Max Sim aims to score better than the random undersampling, while Min Sim does not. This could indicate there are a lot of overlaps between the active and inactive, and therefore, learning the decision boundaries between the two classes is important for the model to distinguish the testing data. On the other hand, Min Sim samples inactive that does not contain active, so this makes the decision boundaries clearer, and it would be more effective if inactive and active are very different from each other, but it could run into problems with tricky points hovering the boundaries of actives and inactives. Therefore, Max Sim was able to do better than Min Sim and random due to the careful selection of diverse and representative points. The low standard deviation in Max Sim can validate the robustness of this method. In this example, Random Forest is more robust than Logistic Regressor and Multilayer Perceptron. The choice of algorithm is critical for modeling structure-activity relationships based on molecular fingerprints. Given the high dimensionality (2048 bits) and non-linearity of the fingerprint data, an ensemble method like Random Forest is expected to be more robust than a linear classifier such as Logistic Regression. Random Forest’s ability to model complex interactions without overfitting, through built-in feature randomization and bagging, makes it well-suited for this task.^30^ This aligns with the findings of Svetnik et al., who demonstrated the superior performance of Random Forest in chemical classification problems.^30^ It also does better or similar to MLP, showing that a complex neural network might not always be necessary for a system like this.

For WDR12, Max Sim achieved higher accuracy than random sampling when using the Random Forest model. However, for Logistic Regression and MLP, the accuracy scores were comparable to the random benchmark. In terms of precision, Random Forest also out-performed the other models and undersampling methods, indicating that the data exhibits complex, nonlinear patterns best captured by an ensemble method. Although MLP is designed to model nonlinear relationships, Random Forest performed more effectively—likely due to its ability to handle both nonlinearities and the high-dimensional nature of molecular fingerprints. A clear trade-off between precision and recall was observed, with Max Sim scoring lowest on recall. This reflects the inherent tension between minimizing false positives and false negatives. Despite the lower recall, Max Sim produced the highest overall scores across most metrics, reaffirming its strength as a sampling method. Interestingly, Logistic Regression achieved the best F1 scores, suggesting that simpler models can still perform competitively when paired with well-chosen sampling techniques. Lastly, ROC AUC scores were relatively consistent across Max Sim, Min Sim, and random sampling, indicating that this metric may be less sensitive to sampling strategies in class-imbalanced settings.

## Conclusion

The results demonstrate that the choice of undersampling technique and classifier significantly impacts model performance, particularly for class-imbalanced datasets and high-dimensional molecular fingerprint data. Across both WDR91 and WDR12 datasets, the Max Sim technique consistently outperformed other sampling strategies, particularly in terms of F1 score and accuracy when paired with the Random Forest classifier. This highlights the importance of selecting diverse and representative samples during undersampling, especially in datasets with overlapping class boundaries.

The trade-off between precision and recall was evident in both datasets, underscoring the need to evaluate multiple metrics rather than relying solely on accuracy. In particular, the F1 score emerged as a more reliable indicator of model effectiveness, especially in imbalanced scenarios where high recall or precision alone may provide a misleading view of performance. Models that incorporated clustering during sampling, such as Max Sim and Min Sim, generally outperformed those that did not, confirming that clustering helps expose the model to a broader and more representative subset of inactives, improving the model’s ability to learn meaningful decision boundaries.

Among classifiers, Random Forest proved to be the most robust, handling both the nonlinear relationships and the high-dimensionality of the input data more effectively than Logistic Regression and MLP. Despite MLP’s capacity to model complex patterns, it did not consistently outperform Random Forest, suggesting that in some domains, simpler ensemble methods may be more effective than deeper neural networks. Overall, the findings emphasize that both sampling strategy and model complexity must be carefully balanced. Techniques like Max Sim, combined with flexible classifiers such as Random Forest, can offer strong and stable performance in the presence of high-dimensional and imbalanced biological datasets.

## Data and Software Availability

The code (including scripts for fingerprint generation, splitting, clustering, undersampling, and model training & evaluation) for the DELight software package is available on GitHub https://github.com/mqcomplab/MDANCE.

## Author Contributions

LC: Data curation, formal analysis, investigation, software, validation, visualization, and writing. RAMQ: Formal analysis, methodology, conceptualization, investigation, software, writing, funding acquisition, supervision, and resources.

## Notes

The authors declare no competing financial interests.

## Acknowledgement

RAMQ and LC thank support from the National Institute of General Medical Sciences of the National Institutes of Health under award number R35GM150620.

